## Supporting Information for "Cognitive control of behavior and hippocampal information processing without medial prefrontal cortex"

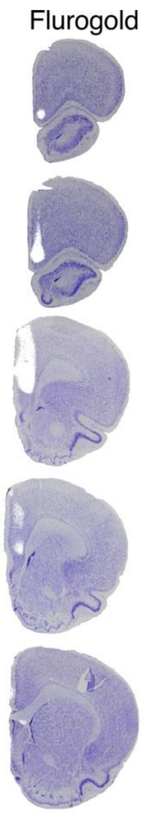

**Figure S1 (related to Figure 1). Targeting the mPFC injections.** To determine the appropriate coordinates for injection of ibotenic acid, we first injected fluorogold and immediately examined the extent of fluorescent labeling, followed by Nissl counterstaining. Fluorogold labeling was confirmed in the cingulate, prelimbic, and infralimbic cortices.

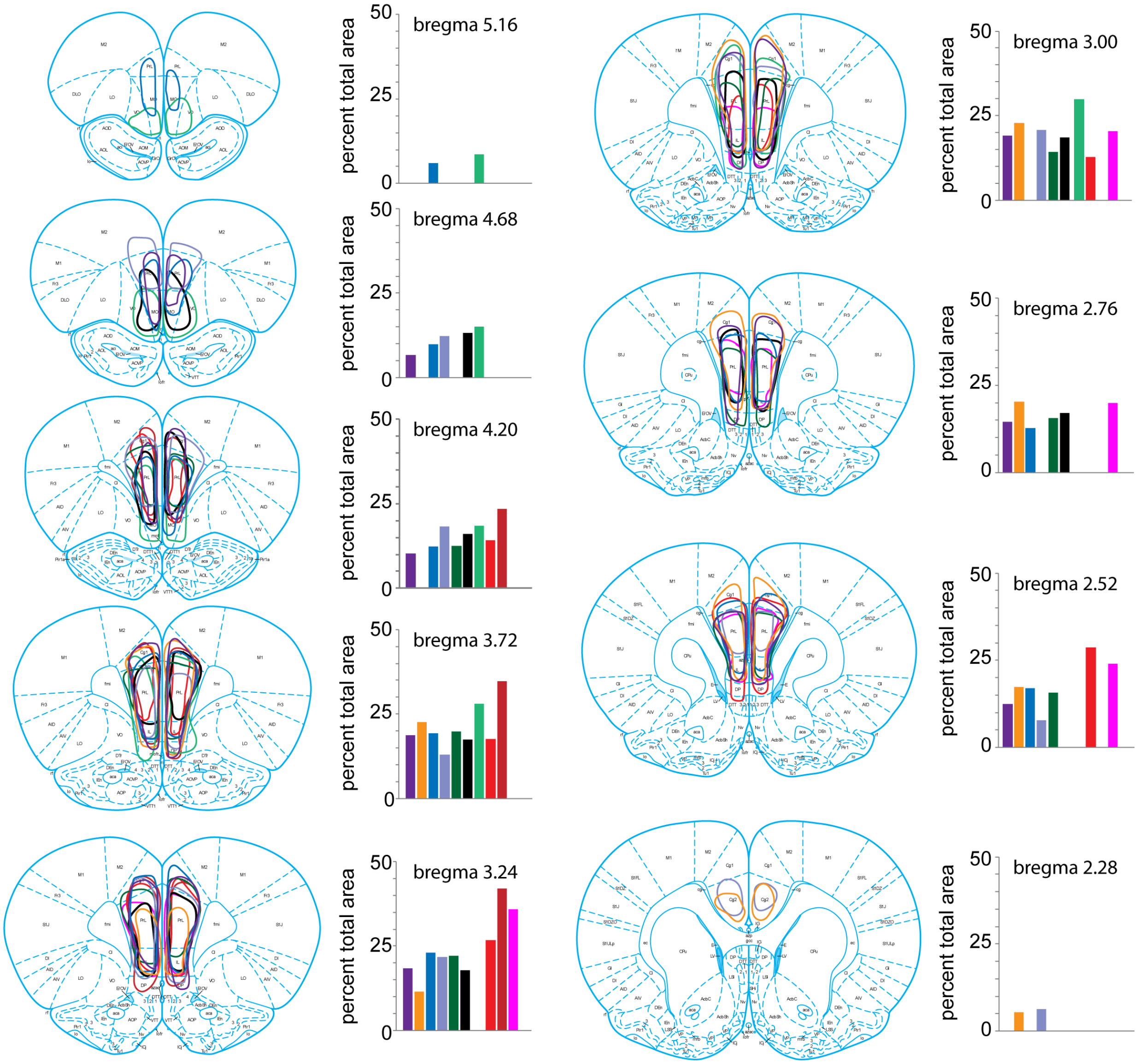

**Figure S2 (related to Figure 1). mPFC lesions.** The extent of each rat's lesion was traced and quantified as the percentage of the total mPFC at each of 9 coronal planes through the A-P extent of the mPFC.

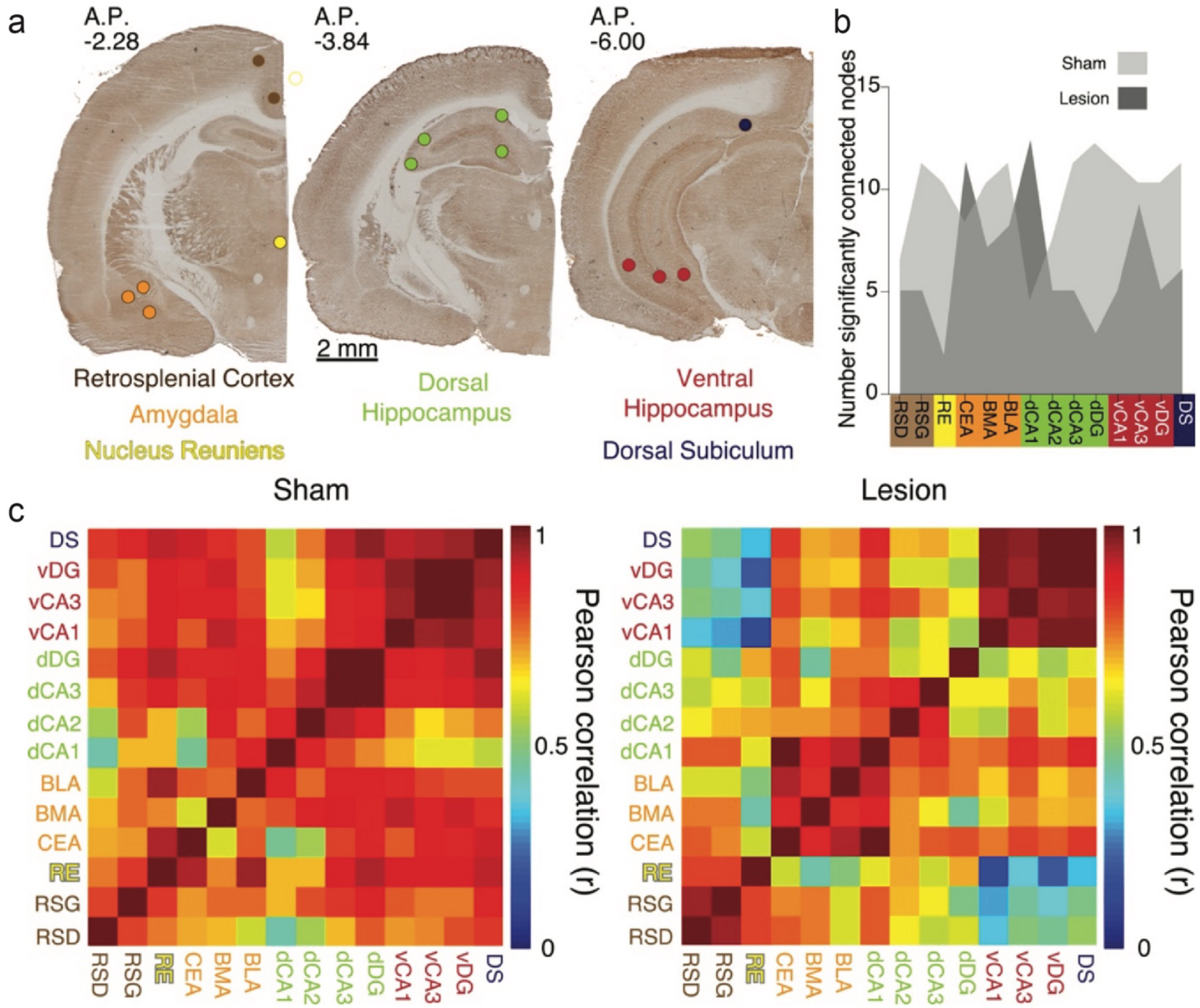

**Figure S3 (related to Figure 2). mPFC lesions change covarying resting-state metabolic activity between the dorsal and ventral hippocampus. a)** Representative cytochrome oxidase staining and locations of the optical density readings. Cytochrome oxidase activity was measured in the dysgranular and granular retrosplenial cortices (RSD and RSG, respectively), the nucleus reuniens (RE), the central nucleus of the amygdala (CEA), basomedial and basolateral amygdala (BMA, and BLA, respectively), the dorsal hippocampus CA1, CA2, CA3 and dentate gyrus areas (dCA1, dCA2, dCA3, dDG, respectively) the ventral hippocampus CA1, CA2 and CA3 areas (vCA1, vCA3, vDG, respectively) and the dorsal subiculum (DS) marked as colored circles. **b)** The matrix of interregional cytochrome oxidase activity correlations indicates that mPFC lesions reduce Pearson correlations in the lesion group. Univariate correlations amongst hippocampus, nucleus reunions, amygdala, and dorsal subiculum decreased after mPFC lesion, but these decreases did not survive the $p < 0.0005$ Bonferroni correction for the 91 comparisons (See Table S1 for abbreviations: BLA-RE: $z = 1.91$, $p = 0.05$, dCA1-CEA: $z = 3.12$, $p = 0.001$, dDG-dCA3: $z = 2.79$, $p = 0.005$, vCA1-RE: $z = 1.94$, $p = 0.05$, DS-RE: $z = 1.92$, $p = 0.05$, DS-dDG: $z = 1.9$, $p = 0.05$). **c)** Number of significantly connected nodes at each site before FDR correction. Sham: $n = 8$; Lesion: $n = 8$.

| Brain Region | Sham (Avg. $\pm$ SEM) | Lesion (Avg. $\pm$ SEM) | t value | *p-value* |
|---|---|---|---|---|
| RSD | 1.84 ± 0.09 | 2.01 ± 0.14 | 1.04 | 0.32 |
| RSG | 1.91 ± 0.09 | 2.16 ± 0.09 | 1.86 | 0.08 |
| RE | 1.47 ± 0.08 | 1.66 ± 0.09 | 1.51 | 0.15 |
| CEA | 1.27 ± 0.18 | 1.94 ± 0.19 | 2.61 | **0.02** |
| BMA | 1.31 ± 0.11 | 1.54 ± 0.20 | 0.99 | 0.34 |
| BLA | 1.33 ± 0.17 | 1.72 ± 0.20 | 1.49 | 0.16 |
| dCA1 | 1.51 ± 0.08 | 1.60 ± 0.04 | 0.97 | 0.35 |
| dCA2 | 1.34 ± 0.08 | 1.43 ± 0.06 | 0.98 | 0.34 |
| dCA3 | 1.50 ± 0.10 | 1.63 ± 0.10 | 0.94 | 0.36 |
| dDG | 1.39 ± 0.08 | 1.50 ± 0.10 | 0.85 | 0.41 |
| vCA1 | 1.54 ± 0.15 | 1.74 ± 0.14 | 1.02 | 0.33 |
| vCA3 | 1.73 ± 0.14 | 1.92 ± 0.15 | 0.94 | 0.36 |
| vDG | 1.75 ± 0.12 | 1.92 ± 0.13 | 0.92 | 0.37 |
| DS | 1.36 ± 0.14 | 1.61 ± 0.19 | 1.03 | 0.32 |

**Table S1. Average cytochrome oxidase activity in sham and lesioned brains** .

Relative Cytochrome Oxidase (CO) activity / μm tissue (x $10^{-1}$). Abbreviations - RSD: dysgranular retrosplenial cortex, RSG: granular retrosplenial cortex, RE: the nucleus reuniens, CEA: the central nucleus of the amygdala, BMA: basomedial amygdala, BLA: basolateral amygdala, dCA1: dorsal CA1, dCA2: dorsal CA2, dCA3: dorsal CA3 dDG: dorsal dentate gyrus, vCA1: ventral CA1, vCA3: ventral CA3, vDG : ventral dentate gyrus, and DS: dorsal subiculum.  (Sham; n=8, Lesion; n=8).

| | | | | | | | | | | | |
|---|---|---|---|---|---|---|---|---|---|---|---|
| | | | | | | | | Time to first enter | | | |
| | | | | | cells | | | Pretrain | | **Initial** | **Retention** |
| mPFC01 | 17 | 17.9 | 315.2 | 419.4 | 139.7 | | 14.5 | 2.6875 | 1 | 8.5625 | |
| mPFCS02 | 70 | 10.3 | 128.4 | 103.7 | 58.5 | | 16 | 6.8125 | 4 | 7.4375 | |
| mPFC3 | 15 | 2.9 | 251.8 | 14.1 | 39.5 | | 16 | 5.875 | 12 | 14.25 | |
| mPFC04 | 15 | 77.3 | 169.7 | 107.1 | 117.6 | | 12 | 5.6875 | 2 | 11.5 | |
| mPFC07 | 24 | 0.8 | 99.9 | 32.9 | 30.0 | | 16 | 10.75 | 6 | 10.75 | |
| mPFC08 | 46 | 29.3 | 135.0 | 84.1 | 349.5 | | 22.5 | 4.375 | 11 | 5.5 | |

**Table S2**. Place cell counts and behavioral measures of the recorded rats.

| **Thalamus (187 cells)** | **Sham lesion** | **mPFC Lesion** | **t(d.f); p values** |
|---|---|---|---|
| Firing rate (AP/s) | 14.81 ± 1.74 | 19.00± 2.16 | 1.51(183); 0.13 |
| Burst ratio | 1.08 ± 0.07 | 0.76 ± 0.08 | 2.91(174); 0.004 |

**Table S3**. Electrophysiological properties of thalamic cells.
